# Influence of amino acids and glucosinolates in three Brassicaceae vegetable plants on preference and performance of the green peach aphid: *Myzus persicae*

**DOI:** 10.1101/2022.05.27.493770

**Authors:** Muhammad Afaq Ahmed, Ning Ban, Sarfaraz Hussain, Raufa Batool, Tong-Xian Liu, Yong-Jun Zhang, He-He Cao

## Abstract

The green peach aphid, *Myzus persicae* (Sulzer) is a generalist pest of various host plants, whose feeding preference and growth performance mainly depends on the quantity and quality of nutrients and defensive metabolites in host plants. Here, we studied the preference and performance of *M. persicae* on three major Brassicaceae vegetables in China and measured nutrient (amino acids) and defensive (glucosinolates) metabolites in these plants. We found that *M. persicae* preferred and performed better on Chinese cabbage than cabbage and radish, which may be due to the relatively higher concentration of amino acids and lower levels of glucosinolates in their leaves. The glucosinolates level in cabbage leaves was ten times higher than the other two plants, while the amino acid concentration in radish was only half of the cabbage or Chinese cabbage. The higher concentration of glucosinolates in cabbage and lower levels of amino acids in radish may account for the poorer preference and growth of *M. persicae* on these two plants. These results suggest that both amino acids and glucosinolates in plants are important determinants of the preference and performance of *M. persicae*, which provide new knowledge for the cultivation and breeding of Brassicaceae vegetables.

## Introduction

Aphids feed mostly on the phloem sap of their host plant, which is high in sucrose and lacking in essential amino acids, resulting in an unbalanced diet [1,2]. Aphids get most of their nitrogen and nutrients from free amino acids in phloem sap, and the concentration and composition of amino acids in phloem are essential indicators of nutritional quality for aphids [1,3]. Previous studies on the pea aphid (*Acyrthosiphon pisum*) and the bird cherry-oat aphid (*Rhopalosiphum padi*) showed that aphids fed on a chemically defined artificial diet with reduced amino acid concentration had lower survival, growth, and fecundity [4,5].

The green peach aphid, *M*yzus *persicae* (Sulzer) (Hemiptera: Aphididae), is an important economical polyphagous pest that infests more than 400 plant species, including Solanaceae, Cruciferae, and Leguminosae families [6]. *M. persicae* causes significant losses of crop yield by direct feeding and transmitting plant viruses [7,8]. Amino acids also play a major role in determining the aphid feeding rate, though sucrose is the main feeding stimulant for most aphid species [2,8,9]. For example, sucrose solutions containing glutamine, methionine, or valine were more attractive to *M. persicae* than sucrose solution alone, and *M. persicae* also preferred to feed and performed better on young cabbage leaves with more amino acids than older leaves [10]. Thus, more amino acids in plants generally enhance aphid feeding preference and growth performance [2,8,10]. During feeding, aphids also consume many phytotoxins such as cardiac glycosides, alkaloids, benzoxazinoids, and glucosinolates, which have toxic characteristics of inhibiting growth (antibiosis) or act as a feeding deterrent (antixenosis) against aphids [11,12].

The main phytotoxins in Brassicaceae plants are glucosinolates and especially their breakdown products, which generally reduce aphid feeding and growth [13,14]. For example, indole glucosinolates were unstable in aqueous solutions and showed strong anti-feeding effects on *M. persicae* [15]. However, *M. persicae* still preferred and performed better on young cabbage leaves that contained a higher level of glucosinolates and amino acids [10]. These findings suggest that the effects of glucosinolates on aphid preference and performance are complex because of aphid adaptation or nutrition compensation. Therefore, future plant-aphid interaction studies should take both nutrients and toxins into account.

In this study, we first investigated the preference and performance of *M. persicae* on three major Brassicaceae crops of China and then monitored the feeding behavior of *M. persicae*. The amino acid, sugar, and glucosinolate levels in the leaves of these three host plants were also measured. This study not only explores the importance of both plant nutrients and defensive metabolites in plant-aphid interactions but also provides new knowledge for the cultivation and breeding of Brassicaceae crops.

## Materials and methods

### Plants and insects

Cabbage (*Brassica oleracea* L. var. ‘Jing Feng 1’), Chinese cabbage (*Brassica rapa pekinensis* var. ‘Dayu’), and radish (*Raphanus sativus* L. var. ‘Weixian K-57’) seeds were sown in the damped garden soil mixture in a controlled plant growth room at (22 ± 2)°Cwith 50% relative humidity and 16:8 h L/D cycle. Plants at three weeks of age with three true leaves were used in all experiments. *Myzus persicae* were obtained from the sweet pepper plants in the field of Qingdao Agricultural University (Qingdao, Shandong, China) and were reared on each host plant for at least three generations in cages under the same conditions.

### Feeding preference of *M. persicae*

Leaf discs (2 cm in diameter) from the three plants were cut and placed equally apart inside one Petri dish (9 cm in diameter) layered with moistened filter paper to maintain the leaf turgor. Fifteen apterous adult aphids were placed in the center of each dish. The number of aphids that had settled on each leaf disc and did not move in 2 min was counted 1, 3, 8, 12, and 24 h after release. Ten replicates were performed for this assay as each dish was a replicate. This assay was replicated three times with aphids from one host plant each time.

### Performance assay

Three apterous adult *M. persicae* from each host plant were confined on the second true leaf of the respective host plant by the nylon mesh bag [8]. Adult *M. persicae* were removed from the leaves after 24 h, leaving five or six newborn nymphs on each leaf. After an additional ten days, the live aphids were counted and adults were weighed on the microbalance (MSA 3.6 P-000-DM, resolution 0.001 mg, Sartorius, Gottingen, Germany). Ten replicates were performed for each host plant.

### Feeding behavior of *M. persicae*

The electrical penetration graph (EPG) direct-current system (Giga-8, Wageningen, Netherlands) was used to record *M. persicae* feeding behavior. A gold wire (18 µm in diameter) was attached to the dorsum of each adult aphid using silver conductive glue [16]. Feeding activities of aphids from respective hosts were recorded for 8 h at 24°C in a Faraday cage. The Stylet+d program was used to record the signal, and the Stylet+a software to label and analyze the EPG waveforms [17]. EPG parameters were calculated automatically by the EPG data 4.3 Excel workbook [18]. Each adult aphid and plant were used only once and were considered as one replicate. Overall, we obtained 15 successful replicates for each host plant.

### Amino acid extraction and analysis

Free amino acids were extracted by grinding 100 mg fresh leaf tissue with a glass mortar and pestle in 0.1 M HCl solution and were analyzed by the LTQ-XL linear ion trap mass spectrometer (LC-MS/MS; Thermo Fisher Scientific, Waltham-MA, USA) [19]. The XTerra MS C18 Column (125 Å pore size, 5 µm, 4.6×150 mm; Waters, Milford, MA, USA) was used for liquid chromatographic separations [20]. The elution of amino acids from the sample solution was conducted in a three-step gradient with mobile phase A (5% acetonitrile and 0.1% formic acid) and mobile phase B (100% acetonitrile). The spray voltage was fixed at 4.5 kV, while the temperature of the ion transfer capillary was set at 320°C. The masses of precursors and productions of each amino acid can be found in our previous work [20]. The external standard amino acid mixture of known concentrations (AAS18, Sigma-Aldrich, St. Louis, MO, USA) was used for quantification, accompanied by cysteine, tryptophan, asparagine, and glutamine [20]. Eight replicates were performed for each host plant.

### Sugar extraction and analysis

The sugars (fructose, glucose, and sucrose) were extracted from plants by grinding 100 mg fresh leaves in 1 ml MilliQ water and were measured by the LTQ-XL linear ion trap mass spectrometer [8]. External standard sugars mixture was used for quantification. Overall, eight replicates were carried out for each host plant.

### Glucosinolate extraction and analysis

Fresh leaf (100 mg) was placed in a 1.5 mL centrifuge tube and kept in the boiling water for 2-3 min to deactivate the myrosinase activity in leaves [21]. The leaves were then ground in MilliQ water (1 mL) with a glass mortar and pestle and the mixture was then centrifuged for 20 min at 12,000 *g*, 4°C. Glucosinolates in the supernatant were measured by the LTQ-XL linear ion trap mass spectrometer [10,21]. The relative concentration of glucosinolates was determined by the standard curve made by 2-propenyl glucosinolate (sinigrin). Eight replicates were performed for each host plant.

### Statistical analysis

The preferences of aphid for different host plant leaves were analyzed independently by paired *t*-test. The equality and normality of the variances were determined using the Kolmogorov–Smirnov and Shapiro-Wilk tests for performance assay, EPG results, individual and total amino acid concentration and glucosinolate concentration in the leaves of different host plants. The ln (1 + x) transformation was used to modify any data that did not match the uniformity of variance assumptions. The effect of different host plant leaves on these factors were then analyzed using one-way analyses of variance (ANOVA) followed by Tukey’s honestly significant difference (HSD) post hoc tests at a significance level of *P* < 0.05. All the above analyses were conducted using R 3.2.2 (R Foundation for Statistical Computing, Vienna, Austria). All figures and graphs were produced in GraphPad Prism v. 8.4.3 for Windows and illustrations were in Adobe Illustrator Package 2021.

## Results

### Preference and performance of *M. persicae*

Significantly more adult aphids reared on cabbage (Fig 1) as well as Chinese cabbage and radish (data not shown) settled on Chinese cabbage leaves compared with cabbage (*t* = 11.59, df = 19, *P* = 0.001; Fig 1A) or radish (*t* = 5.34, df = 19, *P* = 0.001; Fig 1B) plant leaves after release. When cabbage and radish plant leaves were tested, aphids significantly settled on cabbage plant leaves (*t* = 6.24, df = 19, *P* = 0.001; Fig 1C) after release. Aphids feeding on Chinese cabbage resulted in significantly more nymph production (*F* = 28.54, df = 2, 27, *P* < 0.001; Fig 1D) and higher adult body weight (*F* = 6.97, df = 2, 27, *P* < 0.05; Fig 1E) than aphids feeding on radish and cabbage.

**Fig 1.**
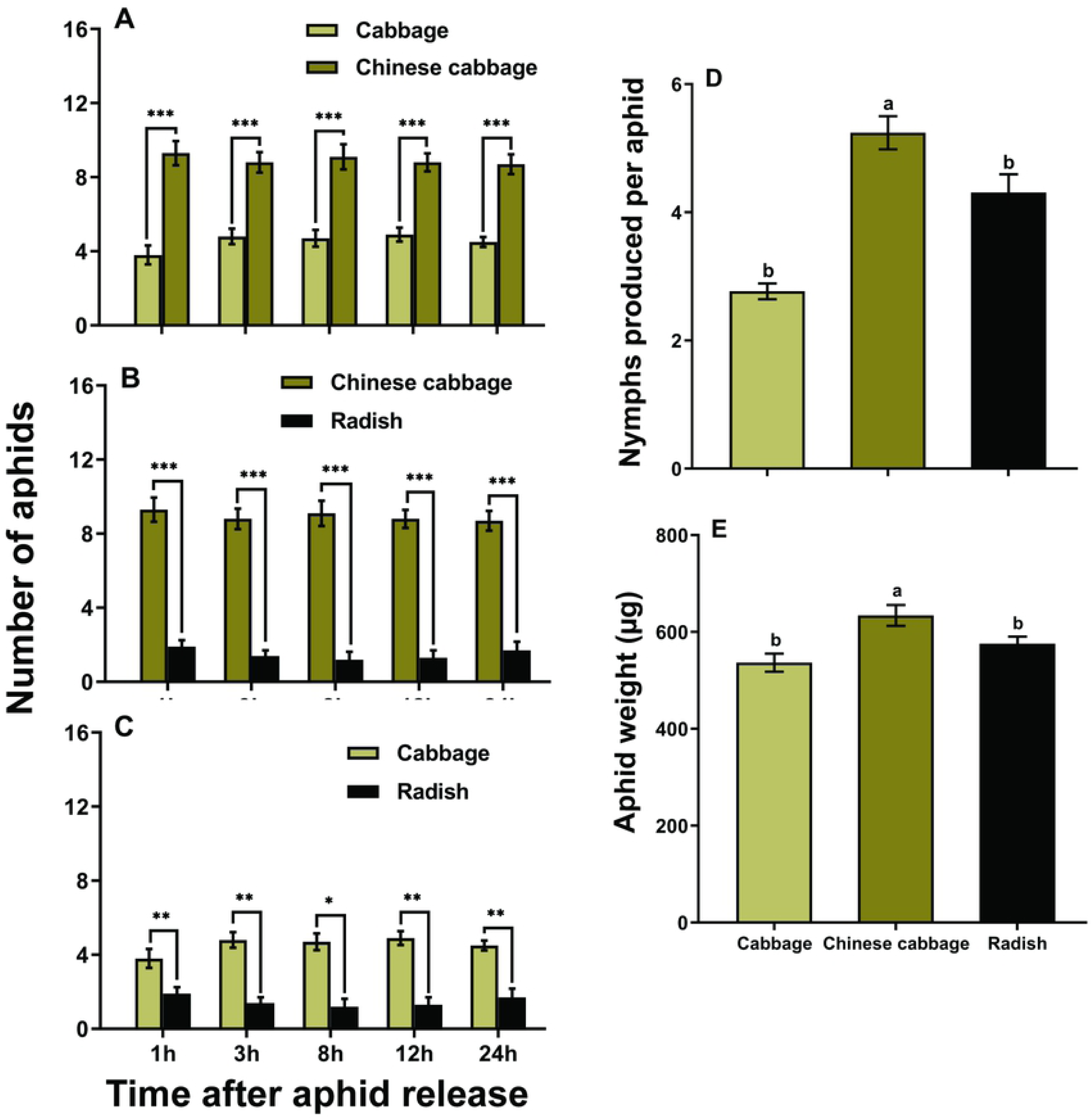
Feeding preference of adult *M. persicae* for different host plant leaves (A-C) (paired *t*-test: * *P* < 0.05, ** *P* < 0.01, *** *P* < 0.001); and growth performance of adult *M. persicae* in terms of nymph production (D); and body weight (E). Values are mean ± SE. Different letters above bars indicate significant differences at each time point or among plants (one-way ANOVA, Tukey’s (HSD) test at *P* < 0.05).

### Aphid feeding behavior

The total probing time of aphids feeding on different host plants was comparative (*P* = 0.957; Table 1). The mean duration of E2 (phloem feeding) (*F* = 5.91, df = 2, 42, *P* < 0.05) and total duration of E2 (*F* = 9.61, df = 2, 42, *P* < 0.05) were significantly shorter when aphids feeding on radish leaves compared with other host plants. No significant difference in the number of E1 (salivation phase) (*P* = 0.194) and E2 (*P* = 0.192) on different host plants was found. Significantly a greater number of probes (*F* = 8.91, df = 2, 42, *P* < 0.001) and short probes (< 3 min) (*F* = 4.32, df = 2, 42, *P* < 0.05) were exhibited by aphids feeding on radish plants (Table 1).

**Table 1.** Probing behavior of *M. persicae* on different host plants.

|  | **Cabbage** | **Chinese cabbage** | **Radish** |
| --- | --- | --- | --- |
| **EPG parameters** | **n = 15** | **n = 15** | **n = 15** |
| Total duration of probing (h) | 7.54 ± 0.23 | 7.67 ± 0.09 | 7.61 ± 0.45 |
| Total duration of E2 (h) | 5.82 ± 0.34 **a** | 4.83 ± 0.62 **a** | 2.61 ± 0.57 **b** |
| Mean duration of E2 (h) | 3.42 ± 0.62 **a** | 1.76 ± 0.56 **b** | 1.01 ± 0.23 **b** |
| Total duration of E1 (min) | 5.13 ± 1.00 | 4.03 ± 0.87 | 3.74 ± 0.68 |
| Mean duration of E1 (min) | 0.94 ± 0.16 **ab** | 1.19 ± 0.15 **a** | 0.76 ± 0.04 **b** |
| Number of E2 | 5.86 ± 1.05 | 3.06 ± 0.95 | 4.53 ± 1.18 |
| Number of E1 | 6.2 ± 1.06 | 3.4 ± 0.94 | 4.7 ± 1.19 |
| Number of probes | 9.0 ± 1.70 **b** | 7.2 ± 1.74 **b** | 19.6 ± 3.02 **a** |
| Number of short probes (< 3 min) | 4.46 ± 1.13 **b** | 3.4 ± 1.08 **b** | 8.8 ± 1.79 **a** |
| Number of probes to the first E1 | 4.46 ± 1.14 | 6.73 ± 1.57 | 7.53 ± 1.86 |
| Duration of np period before the first E1 (h) | 0.07 ± 0.02 | 0.20 ± 0.08 | 0.18 ± 0.06 |
| Duration of first probe (min) | 119.30 ± 47.95 | 49.40 ± 31.16 | 25.92 ± 7.82 |
| Time from first probe to first E1 (h) | 0.73 ± 0.11 | 1.29 ± 0.24 | 1.20 ± 0.24 |

Data are means ± SE and different letters within each row indicate significant differences (one-way ANOVA, Tukey's (HSD) test at $P < 0.05$ ). E1, salivation; E2, phloem ingestion; np, non-probe; n, number of replicates.

### Amino acid profiles in leaves

The concentration of the essential amino acid threonine (*F* = 45.34, df = 2, 21, *P* < 0.001), and aspartate (*F* = 17.78, df = 2, 21, *P* < 0.001) in cabbage leaves were significantly higher than Chinese cabbage and radish leaves (Fig 2A). Chinese cabbage leaves contained significantly higher levels of asparagine (*F* = 16.46, df = 2, 21, *P* < 0.001), valine (*F* = 19.67, df = 2, 21, *P* < 0.001), isoleucine (*F* = 21.51, df = 2, 21, *P* < 0.001), and phenylalanine (*F* = 23.27, df = 2, 21, *P* < 0.001) (Fig 2A and 2B). Total amino acid concentration in cabbage or Chinese cabbage leaves was also significantly higher than that in radish leaves (*F* = 31.0, df = 2, 21, *P* < 0.001; Fig. 2C).

**Fig 2.**
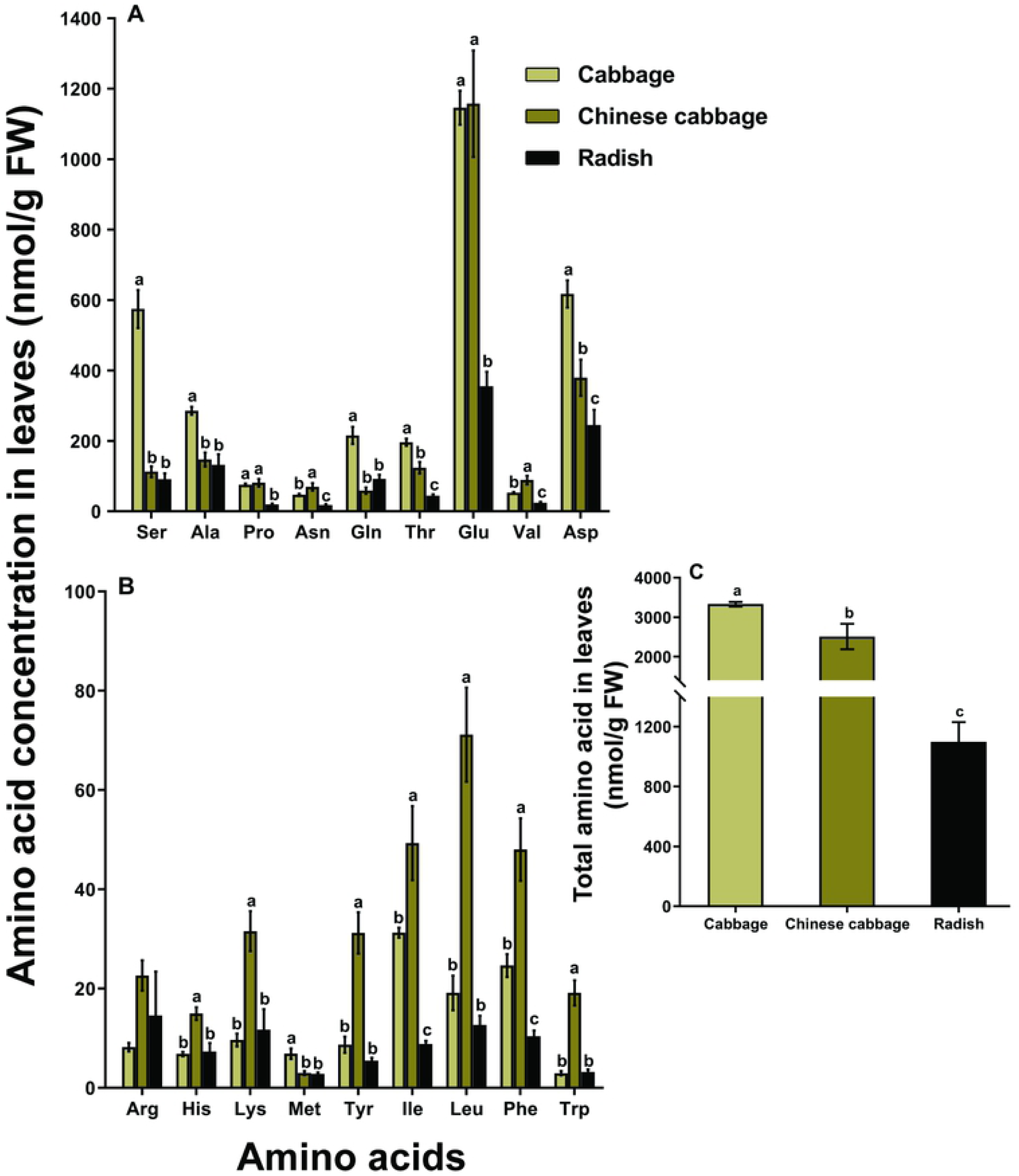
Nutrient quality in the leaves of different host plants. The individual concentration of amino acids (A-B), and the total concentration of amino acids (C). Different letters above bars of each amino acid indicate significant differences among plants (one-way ANOVA, Tukey’s (HSD) test at *P* < 0.05).

### Sugar content in leaves

The concentration of fructose (*F* = 13.38, df = 2, 21, *P* < 0.001; Fig 3A), glucose (*F* = 9.51, df = 2, 21, *P* < 0.001; Fig 3B), sucrose (*F* = 4.01, df = 2, 21, *P* < 0.05; Fig 3C), and total sugar (*F* = 12.0, df = 2, 21, *P* < 0.001; Fig. 3D) in cabbage leaves were significantly higher than in Chinese cabbage and radish leaves.

**Fig 3.**
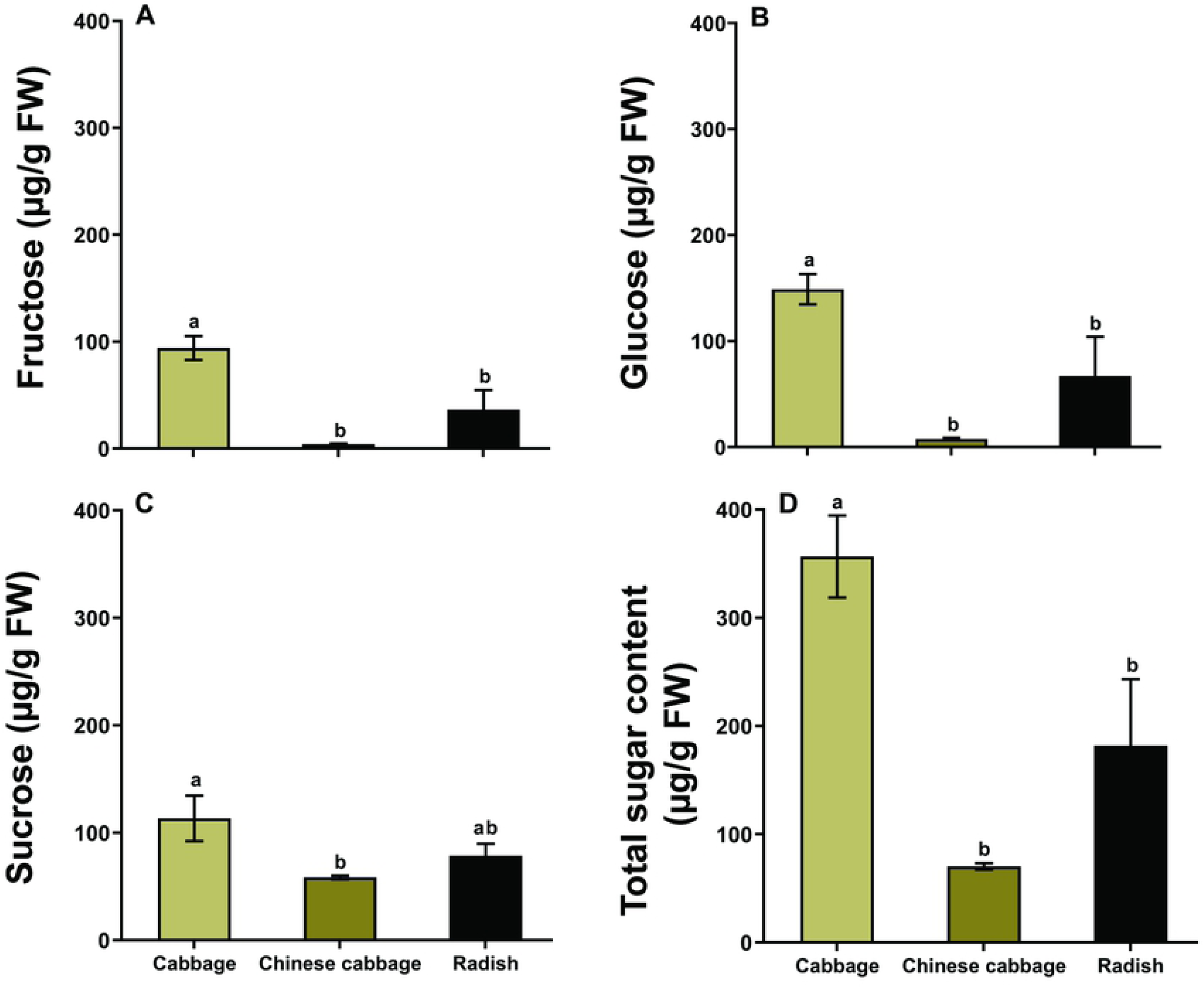
Sugar content in the leaves of different host plants. fructose (A); glucose (B); sucrose (C), and total sugar content (D). Different letters above bars indicate significant differences (one-way ANOVA, Tukey’s (HSD) test at *P* < 0.05).

### Glucosinolate concentration in leaves

The concentrations of glucoiberin (GIB), glucoraphanin (GRA), and sinigrin (SIN) were the highest in cabbage, while these glucosinolates were undetectable in Chinese cabbage and radish leaves. However, the level of progoitrin (PRO) was significantly higher in Chinese cabbage leaves (*F* = 12.05, df = 2, 21, *P* < 0.05; Fig 4A). Indolic glucosinolates 4-hydroxyglucobrassicin (4HGBS), glucobrassicin (GBS), and 4-methoxyglucobrassicin (4MGBS) were found in all three hosts. The concentration of GBS was significantly higher in cabbage leaves (*F* = 59.9, df = 2, 21, *P* < 0.001; Fig 4A), while the concentration of 4MGBS was the highest in radish leaves (*F* = 16.21, df = 2, 21, *P* < 0.001; Fig 4A). Neoglucobrassicin (NGBS) with the highest concentration was found in the Chinese cabbage leaves (Fig 4A). The total content of glucosinolates in cabbage leaves was about ten times and 40 times higher than Chinese cabbage and radish leaves, respectively (*F* = 140.0, df = 2, 21, *P* < 0.001; Fig 4B).

**Fig 4.**
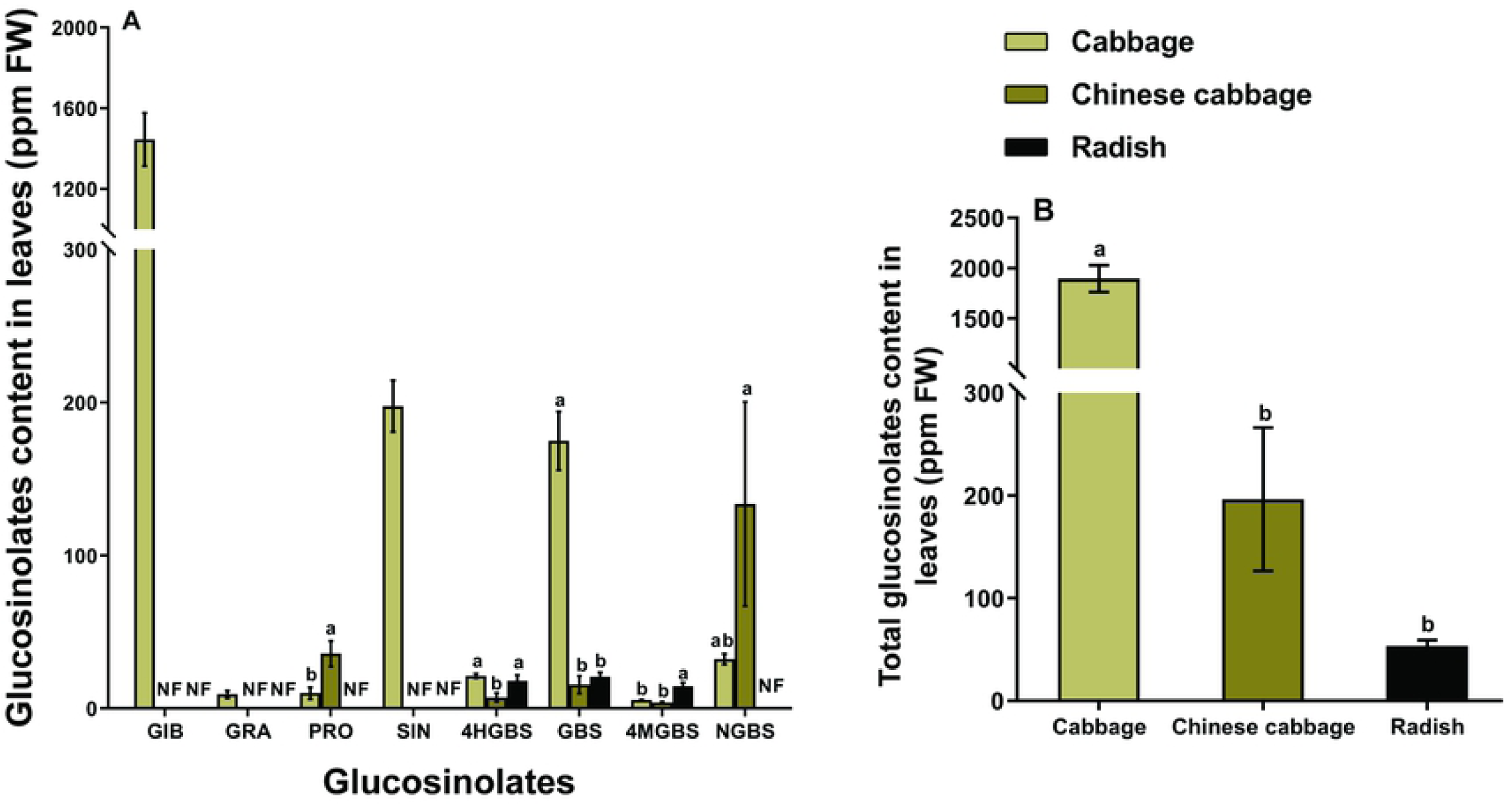
The concentration of glucosinolates in the leaves of different host plants. Individual concentration of glucosinolates (A), and total concentration of glucosinolates (B). Different letters above bars indicate that each glucosinolate is significantly different among plants (one-way ANOVA, Tukey’s (HSD) test at *P* < 0.05). Glucosinolate side-chain abbreviations: GIB = glucoiberin; GRA = glucoraphanin; PRO = progoitrin; SIN = sinigrin; 4HGBS = 4-hydroxyglucobrassicin; GBS = glucobrassicin; 4MGBS = 4-methoxyglucobrassicin; NGBS = neoglucobrassicin.

## Discussion

Our results showed that *M. persicae* preferred and performed better on Chinese cabbage, which may be due to the higher level of nutrient (amino acids) and lower level of glucosinolates in the Chinese cabbage leaves. In contrast, the higher concentration of glucosinolates in cabbage and lower level of amino acids in radish may account for the poorer preference and growth of *M. persicae* on these two plants, respectively. These results suggest that both amino acids and glucosinolates in Brassicaceae plants influence the preference and performance of *M. persicae* which is in accordance with previous publications [10,22,23]. In most cases, aphids make their decision to accept the host plant before their stylet reaches phloem sieve element, relying on nutrients or toxins present in the peripheral (non-vascular) plant cells [24]. In the feeding preference assay, significantly more aphids settled on Chinese cabbage leaves since 1 h after release; however, EPG data showed that *M. persicae* took more than 1 h to reach the phloem (E1), indicating that factors that influence aphid feeding host preference may be located at epidermal or mesophyll cells [24].

Compared with cabbage, Chinese cabbage leaves contained relatively lower levels of feeding stimulants (sucrose) and comparable amino acid; thus, more aphids selected Chinese cabbage possibly because of the lower glucosinolate contents in Chinese cabbage leaves. Indole glucosinolates has been shown strong antixenosis effects against *M. persicae*, and we found that the indole glucosinolates 4HGBS and GBS in Chinese cabbage leaves were lower than those in cabbage leaves (Fig. 4A) [13,15]. However, *M. persicae* fed on cabbage showed a longer mean phloem sap feeding time (Table 1) than on Chinese cabbage, which could be explained by the fact that the higher level of sucrose in phloem may mask the taste of glucosinolate. In addition, the leaves of Chinese cabbage plants were soft and thus may have flexible cell walls, allowing aphid stylets to penetrate more easily [25].

Amino acids and glucosinolates are the key nutrition and defensive metabolites respectively, in the Brassicaceae plants that determine aphid performance [3,26,27]. In this study, there was no significant difference in amino acid content between cabbage and Chinese cabbage. In addition, although the mean phloem feeding time was significantly different between Chinese cabbage and cabbage, the total time was not, suggesting that aphids obtain a similar amount of nutrition from these two plants. Thus, the significantly higher level of glucosinolates in cabbage leaves may account for the lower body weight and fecundity of *M. persicae* on this plant. However, in some previous studies, *M. persicae* grew better on young cabbage leaves or pre-infested Chinese cabbage leaves that contain a higher level of glucosinolates, which could be explained by that aphids also ingested considerably more nutrition (mainly the amino acids) from these leaves [8, 10]. Although radish contained a lower level of glucosinolates, it also had fewer nutrients, which may be the reason why aphids had poorer growth and preference on this plant. In addition, aphids had a shorter phloem sap time on radish and thus ingested fewer nutrients when feeding on radish. These results suggest that the effect of glucosinolates in plant resistance against aphids is highly variable, which could be affected by aphid’s detoxification ability or nutrient compensation [6,10]. Nevertheless, we cannot exclude the possibility that some other unknown factors may also be involved in this plant-aphid interaction.

This study reveals that *M. persicae* prefers and performs better on Chinese cabbage, which may be due to both the nutrients and defensive metabolites in the host plants. Our results suggest that Chinese cabbage is more susceptible to *M. persicae* and may need more field monitoring and control measures to manage this pest, and will also provide new knowledge for breeding aphid-resistant Brassicaceae crops.

## Acknowledgments

We are grateful for the assistance of Zhan-feng Zhang (College of Plant Protection, Northwest A&F University, China) for LC/MS/MS analysis and Professor Michael Keller (University of Adelaide, Australia) for revising this manuscript.

## Author Contributions

**Conceptualization:** Muhammad Afaq Ahmed, He-He Cao

**Investigation:** Muhammad Afaq Ahmed, Ning Ban, Sarfaraz Hussain, Raufa Batool

**Methodology:** Muhammad Afaq Ahmed, He-He Cao

**Data curation:** Muhammad Afaq Ahmed

**Validation:** Muhammad Afaq Ahmed, He-He Cao

**Visualization:** Muhammad Afaq Ahmed, He-He Cao

**Supervision:** Muhammad Afaq Ahmed, He-He Cao

**Funding acquisition:** Yong-Jun Zhang, He-He Cao

**Writing - original draft:** Muhammad Afaq Ahmed, He-He Cao

**Writing – review & Editing:** Muhammad Afaq Ahmed, He-He Cao, Tong-Xian Liu, Yong-Jun Zhang

## Notes

### Competing Interest Statement

The authors have declared no competing interest.

